# A high coverage Mesolithic aurochs genome and effective leveraging of ancient cattle genomes using whole genome imputation

**DOI:** 10.1101/2024.01.23.576850

**Authors:** Jolijn A.M Erven, Amelie Scheu, Marta Pereira Verdugo, Lara Cassidy, Ningbo Chen, Birgit Gehlen, Martin Street, Ole Madsen, Victoria E Mullin

**Affiliations:** Groningen Institute of Archaeology, University of Groningen, Groningen, Netherlands; Smurfit Institute of Genetics, Trinity College Dublin, Dublin D02 PN40, Ireland; Palaeogenetics Group, Institute of Organismic and Molecular Evolution (iOME), Johannes Gutenberg-University Mainz, 55099 Mainz, Germany; Key Laboratory of Animal Genetics, Breeding and Reproduction of Shaanxi Province, College of Animal Science and Technology, Northwest A&F University, Yangling 712100, China; Institute for Prehistory and Protohistory, University of Cologne, Weyertal 125, 50931, Cologne, Germany; MONREPOS, Archaeological Research Centre and Museum for Human Behavioural Evolution,Schloss Monrepos, D - 56567 Neuwied, Germany; Animal Breeding and Genomics, Wageningen University and Research, Wageningen, Netherlands

## Abstract

Ancient genomic analyses are often restricted to utilising pseudo-haploid data due to low genome coverage. Leveraging low coverage data by imputation to calculate phased diploid genotypes that enable haplotype-based interrogation and SNP calling at unsequenced positions is highly desirable. This has not been investigated for ancient cattle genomes despite these being compelling subjects for archaeological, evolutionary and economic reasons. Here we test this approach by sequencing a Mesolithic European aurochs (18.49x; 9852-9376 calBC), an Early Medieval European cow (18.69x; 427-580 calAD), and combine these with published individuals; two ancient and three modern. We downsample these genomes (0.25x, 0.5x, 1.0x, 2.0x) and impute diploid genotypes, utilising a reference panel of 171 published modern cattle genomes that we curated for 21.7 million (Mn) phased single-nucleotide polymorphisms (SNPs). We recover high densities of correct calls with an accuracy of >99.1% at variant sites for the lowest downsample depth of 0.25x, increasing to >99.5% for 2.0x (transversions only, minor allele frequency (MAF) ≥2.5%). The recovery of SNPs correlates with coverage, on average 58% of sites are recovered for 0.25x increasing to 87% for 2.0x, utilising an average of 3.5 million (Mn) transversions (MAF ≥2.5%), even in the aurochs which is temporally and morphologically distinct from the reference panel. Our imputed genomes behave similarly to directly called data in allele-frequency-based analyses; for example consistently identifying runs of homozygosity >2mb, including a long homozygous region in the Mesolithic European aurochs.

## Introduction

Cattle are ubiquitous and have been a significant economic and cultural resource for millennia. Taurine cattle (*Bos taurus*) were initially domesticated from the now extinct aurochs (*Bos primigenius*) *circa* 10,500 years BP in southwest Asia (Vigne et al., 2017), with additional introgression from local wild populations in Africa, Europe and the Levant (Park et al., 2015; Verdugo et al., 2019). Indicine cattle (*Bos indicus*) are adapted to warmer climates and descend from the recruitment of a divergent aurochs population (*Bos namadicus*), likely in the Indus Valley region ∼8,000 years BP (Patel & Meadow, 2017). Cattle genetic variation is highly studied but their evolutionary history is incompletely understood, with ancient genome investigation required to uncover key processes in prehistory. For example, human-mediated movement of indicine cattle resulted in widespread admixture between taurine and indicine cattle in southwest Asia at least 4,200 years BP, resulting in hybrid cattle which persist in the region today (Verdugo et al., 2019). However, as archaeological remains are usually low in overall DNA concentration and endogenous DNA content, ancient genomes are typically low coverage (<5x) (Daly et al., 2018; Frantz et al., 2019; Librado et al., 2021; Verdugo et al., 2019), preventing confident diploid calling and limiting analyses to pseudo-haploid data, the sampling of one allele per variant site.

Genotype imputation - the statistical inference of unobserved genotypes by utilisting reference panels of haplotypes - is now a widely used methodological approach in ancient human genomics (Cassidy et al., 2020; Gamba et al., 2014; Martiniano et al., 2017). Specifically the development of GLIMPSE, a tool created for imputation from low-coverage genomes (Rubinacci et al., 2021), has enabled efficient imputation of large ancient human datasets (Clemente et al., 2021; Hui et al., 2020; Rohland et al., 2022; Sousa da Mota et al., 2023)ancient pigs (Erven et al., 2022). Imputing low coverage ancient genomes enables inferences of genome-wide diploid genotypes, diversifying analyses to include haplotype-focused or genealogical methods; *e.g*. inference of autozygosity within, and identity by descent between genomes. While genotype imputation is regularly used in modern livestock genetics, including cattle, (Hayes & Daetwyler, 2019), the efficacy of imputation has not been explored in ancient cattle.

In order to explore the imputation and subsequent analyses of ancient *Bos* we sequenced high coverage genomes (>18x) of a Mesolithic German female aurochs and an Early Middle Ages Dutch cow. We combine these with two previously published high (>13×) coverage ancient genomes (a Neolithic Anatolian domesticate and a Medieval taurine x indicine hybrid from Iraq (Verdugo et al., 2019) and three modern cattle genomes of different ancestries (European *Bos taurus*, African *Bos taurus* and *Bos indicus*). A reference panel of 171 publicly available modern cattle genomes, composed of *Bos taurus* and *Bos indicus*, was aligned and curated for 21.7 million (Mn) single-nucleotide polymorphisms (SNPs). By downsampling the test individuals, imputing with GLIMPSE and comparing analyses from direct genotype calls we establish that this approach is both feasible and desirable for leveraging low coverage ancient genome sequences, even for the extinct European aurochs.

## Results & Discussion

### The first high coverage ancient European aurochs and domestic cattle genomes

Two ancient cattle genomes have been published with sequencing density sufficient for accurate genome-wide genotype calls; a Neolithic Turkish sample (Sub1; 6221-6024 calBC, 13.85×) and a Medieval taurus-indicus hybrid sample from Iraq (Bes2; 1295-1398 calAD, 13.8x) (Verdugo et al., 2019). Here we additionally report the first ancient European domesticate to high coverage, a 18.69x Early Middle Ages Dutch female (Win1) from Winsum-Bruggeburen (424-569 calAD). We also report the first high coverage Mesolithic female European aurochs (Bed3; 9852-9376 calBC) 18.49x genome excavated in Bedburg-Königshoven, Germany (Table SI 1). When we apply standard allele-frequency-based analysis methods (Figure 1) Win1 clusters with modern southern and central European cattle and Bed3 falls close to a published younger Mesolithic British aurochs (CPC98 - 5746-5484 calBC (Park et al., 2015). As expected, Sub1 clusters with other Neolithic Anatolian samples, while the position of Bes2 confers its hybrid ancestry (Verdugo et al., 2019).

**Figure 1).**
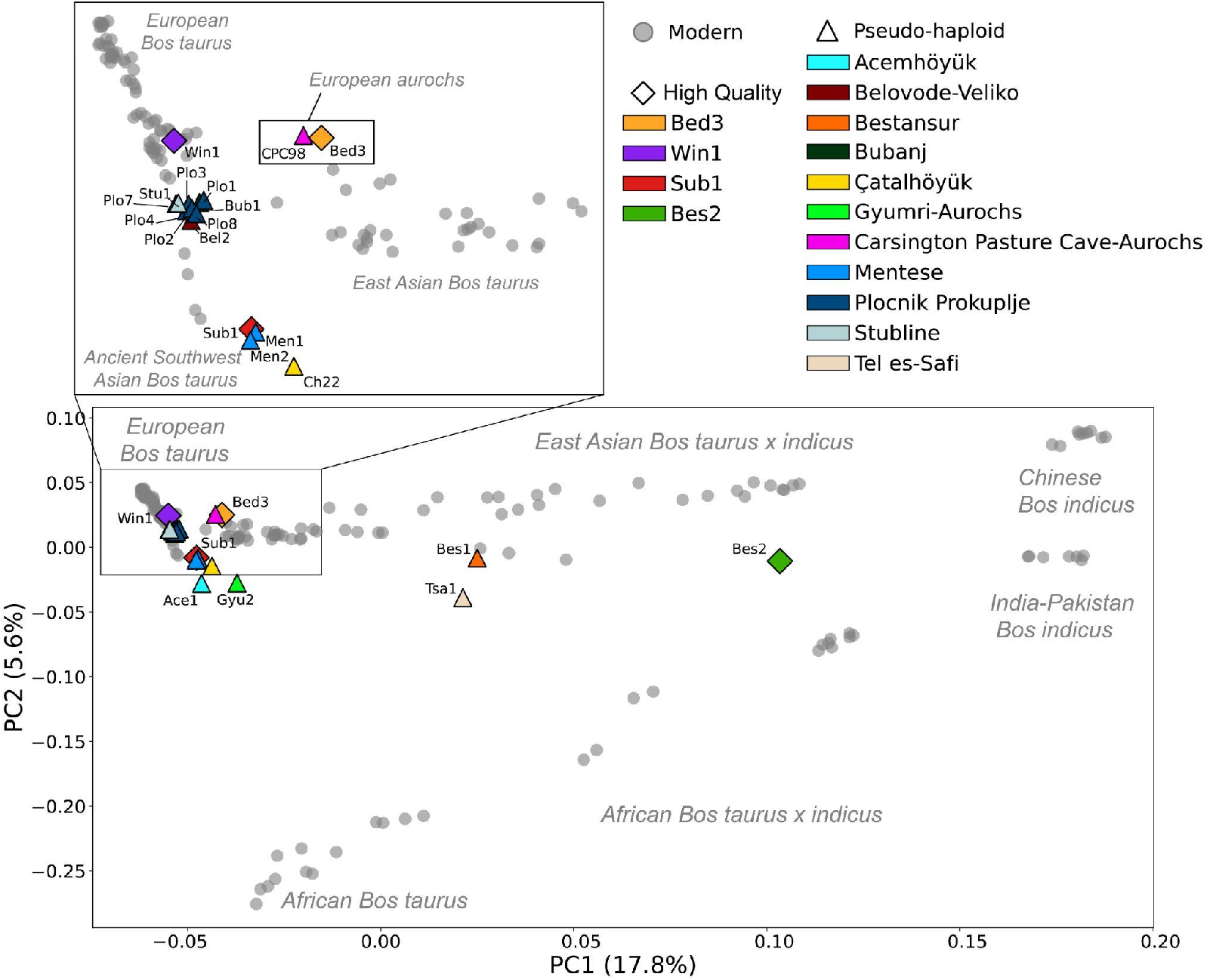
Principal Components analysis of ancient cattle. Bed3 and Win1 along with previously published ancient cattle are projected onto modern cattle diversity shown in grey circles. The four test ancient genomes are represented by diploid calls denoted as diamonds and coloured by sample, while previously published low coverage ancient samples are pseudo-haploid and are denoted by triangles and coloured by site (Table SI 1). Bed3 clusters with the previously published British Aurochs CPC98, while Win1 clusters with modern Southern European *Bos taurus*.

### Ancient cattle genomes impute non-rare alleles to high accuracy

We assembled, aligned and variant called, a phased reference panel of 171 published high coverage (>7.6×) modern cattle genomes of varying ancestries, and with a geographical distribution including Europe, Africa and Asia (Method Section–Variant Discovery; Figure 1, Table SI 2). Variants called with Graphtyper were curated with stringent filters (Methods-Variant Discovery), resulting in 21,656,052 high-confidence SNPs, including 6,521,311 transversions (Table SI 2).

The four ancient *Bos* along with three previously published modern genomes (European *Bos taurus,* African *Bos taurus* and *Bos indicus*) were successfully imputed from a range of downsampled coverages (0.25×, 0.5×, 1×, 2×) using the newly created reference panel (Method section–Variant Discovery). Imputation accuracy, the concordance between imputed genotypes (GP ≥0.99 and INFO ≥0.99) and high quality validation genotypes for the high coverage genomes was calculated for heterozygous and homozygous alternative sites via PICARD GenotypeConcordance (“Picard Toolkit,” 2019); this accuracy is relatively stable over the different genome coverages when tested with transversions only (Figure 2a; Table SI 3). However, rare alleles are affected more by a reduction in genome coverage (Figure 2a). Notably, this trend is also present when all sites (transitions and transversions) are considered (Figure SI 1; Table SI 4).

**Figure 2).**
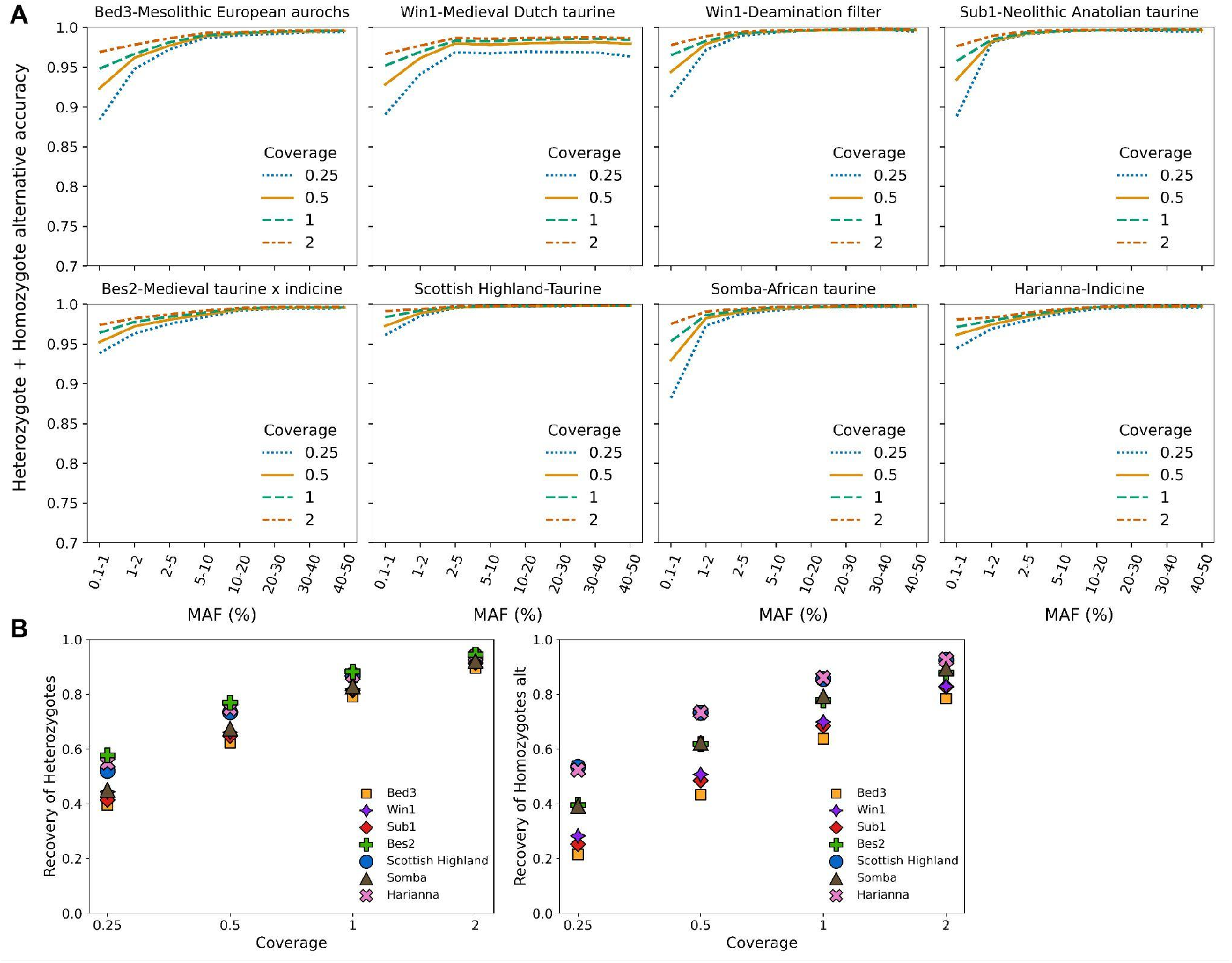
**A**) Accuracy of imputation of heterozygous and homozygous alternative sites at different minor allele frequency (MAF) bins and different downsampled coverages, transversions only and with a GP and INFO filter ≥0.99. Downsampled coverage is denoted by line style and colour. An additional graph is included for Win1 to demonstrate the positive effect on the accuracy of a deamination filter prior to imputation. **B**) Recovery rate of heterozygotes and homozygotes alternative for the different downsampled coverages (MAF ≥ 2.5%, transversions only, GP ≥0.99 and INFO ≥0.99), with samples denoted by shape and colour.

Across the coverage range, more common alleles have high accuracy, but this reduces for rarer alleles, with a marked falling off at minor allele frequency (MAF) <5% (Figure 2a). The lowest rare allele accuracies are observed in the Mesolithic European aurochs, Neolithic Anatolian domestic and modern African taurine samples. This reduction in accuracy is likely due to the underrepresentation of these ancestries in the reference panel (Method section– Variant Discovery; Table SI 2). A similar trend is also observed in non-European ancient humans (Cassidy et al., 2016; Gamba et al., 2014; Martiniano et al., 2017; Sousa da Mota et al., 2023). Additionally, heterozygous sites have a higher accuracy than homozygous alternative sites at more common (MAF >10%) alleles (Table SI 3).

When we consider transversions with a MAF minimum threshold of 2.5%, the lowest accuracy is observed at the lowest downsample coverage of 0.25× (Win1 96.8%, Bed3 99.08%, Bes2 99.1%, Sub1 99.52%) (Table SI 5). It is interesting that, despite the highest temporal (>10,000 yr) distance from the contemporary reference panel, the aurochs Bed3 performs well, for example better than the more recent European domesticate Win1. This implies that the modern reference imputation panel contains a substantial degree of segregating European wild haplotype variants, presumably due to introgressions over the thousands of years when wild and herded animals cohabited on the continent (Park et al., 2015; Verdugo et al., 2019). A similar pattern has also been observed in the imputation of ancient humans, when Indigenous Americans are accurately imputed despite the lack of unadmixed Indigenous American genomes in a reference panel (Cassidy et al., 2016; Gamba et al., 2014; Martiniano et al., 2017; Sousa da Mota et al., 2023). In our data, despite its relatively young provenance, Win1 has an elevated damage profile at CpG sites in the first and last 30bp of the sequencing reads (Figure SI 2). When we filter potential deamination signals in this genome prior to imputation (Figure 2a, Figure SI 3) we demonstrate an improvement in imputation accuracy at each downsampled coverage; at the lowest depth of 0.25× accuracy increases from 96.8% to 99.6% for transversions with MAF ≥ 2.5%.

Our ancient samples are of diverse provenance. In addition to the European wild and northwest European domestic genomes, we also successfully impute an Anatolian Neolithic *Bos taurus* proximal to the origins of cattle in Southwest Asia and a *Bos indicus x Bos taurus* hybrid from Iraq. These results suggest a wide potential for accurate genotype imputation of ancient cattle.

### Ancient imputation achieves genome-wide genotype recovery

Variation is demonstrated in the proportion of genotypes which are recovered, with a clear positive trend between genome coverage and site recovery rate (Figure 2B & Table SI 3-5). The average recovery rate of heterozygote, homozygote alternative and homozygote reference sites for the ancient genomes at a depth of 2× was 87.3% (3.0 Mn sites), while a rate of 58.0% (2.1 Mn sites) was achieved at a depth of 0.25× (MAF ≥2.5%, transversions only, GP ≥0.99 and INFO ≥0.99) (Table SI 5). Across the downsampled coverages the recovery rate for heterozygote and homozygote alternative sites was higher in the modern genomes (39.2%-92.7% at 0.25×-2× coverage) than the ancients (30.6%-91.2%) (MAF ≥2.5%, transversions only) (Figure 2B). Additionally, when partitioning the data by MAF bins the recovery rate differs between heterozygotes and homozygotes alternative to the reference genome, where heterozygotes have a higher rate with more common alleles than homozygotes alternative, a trend which reverses with rare alleles, mirroring the accuracy results (Table SI 3).

### Imputed genomes allow accurate analysis outcomes

We conducted unsupervised frequency-based analyses which are commonly used in ancient population genomics (*i.e.* PCA, ADMIXTURE) demonstrating a positive trend between accuracy and increasing downsample coverage (Figure SI 5-9). Here we report analyses utilising the 0.5× downsample imputation (Figure 3A). These analyses were accurate; the projection PCA demonstrates clustering between each high-quality and imputed replicate with the greatest eigenvector difference observed for the modern African taurine individual (Somba) across PC2 (Figure 3). ADMIXTURE analysis also estimated similar spectra of ancestral component profiles between imputed and high-quality genotypes (Figure SI 8-9). This successful replication included analysis of the most temporally distant sample, the ∼11,500 yr old Mesolithic European aurochs (Bed3).

**Figure 3).**
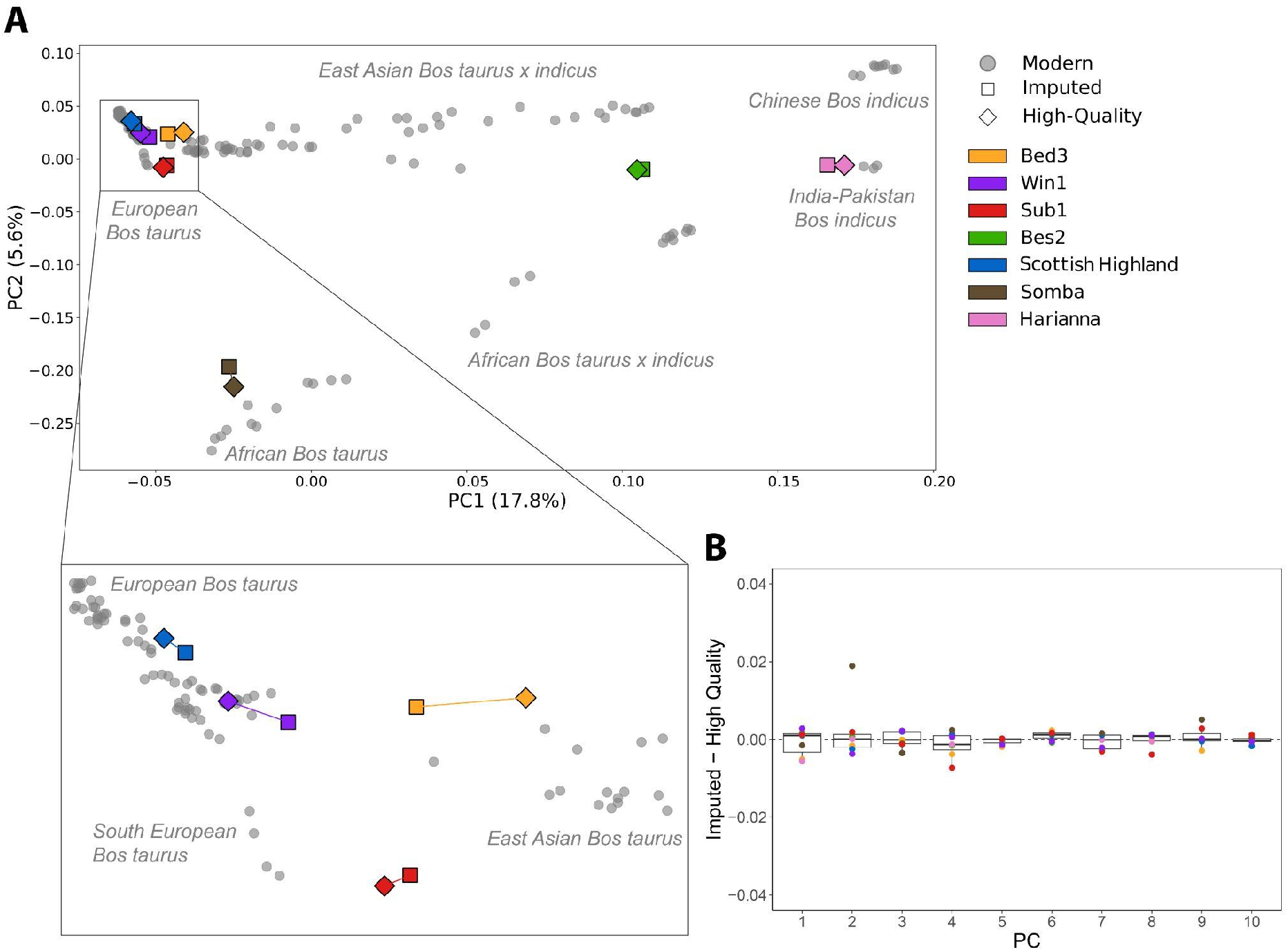
Principal Components Analysis (PCA) of imputed and high-quality genotypes of the seven test individuals onto the modern cattle reference panel. **A:** Projections for 0.5× imputed and high quality diploid genotypes of the test samples onto the modern reference panel along the first two eigenvectors. Dataset was filtered for a MAF ≥2.5%, transversions and LD pruning resulting in 812,801 SNPs. Test individuals are denoted by colour with imputed and high-quality represented by squares and diamonds respectively, while the reference panel individuals are plotted as grey circles. **B:** Boxplots of the normalised differences in the coordinates of the high-quality and 0.5× imputed genomes for the first ten principle components. The horizontal lines of the box pot represent the first quartile, median and third quartile, whiskers represent 1.5 times the quartile range.

For the first time we applied a runs of homozygosity (ROH) analysis on ancient cattle and compared high-quality and imputed data (Figure 4; Figure SI 10). Patterns of ROH were consistent when comparing the 0.5× downsampled imputation to the high-quality genotypes (Figure 4; Figure SI 11). This was true for both genome-wide summaries of ROH (Figure 4A, B) and in the detail of specific genome locations (Figure 4C; Figure SI 11). For example, a large ROH of 15.8 Mb on chromosome 12 of the Mesolithic aurochs is identified using both imputed and high-quality genotypes (Figure 4C; Figure SI 12).

**Figure 4).**
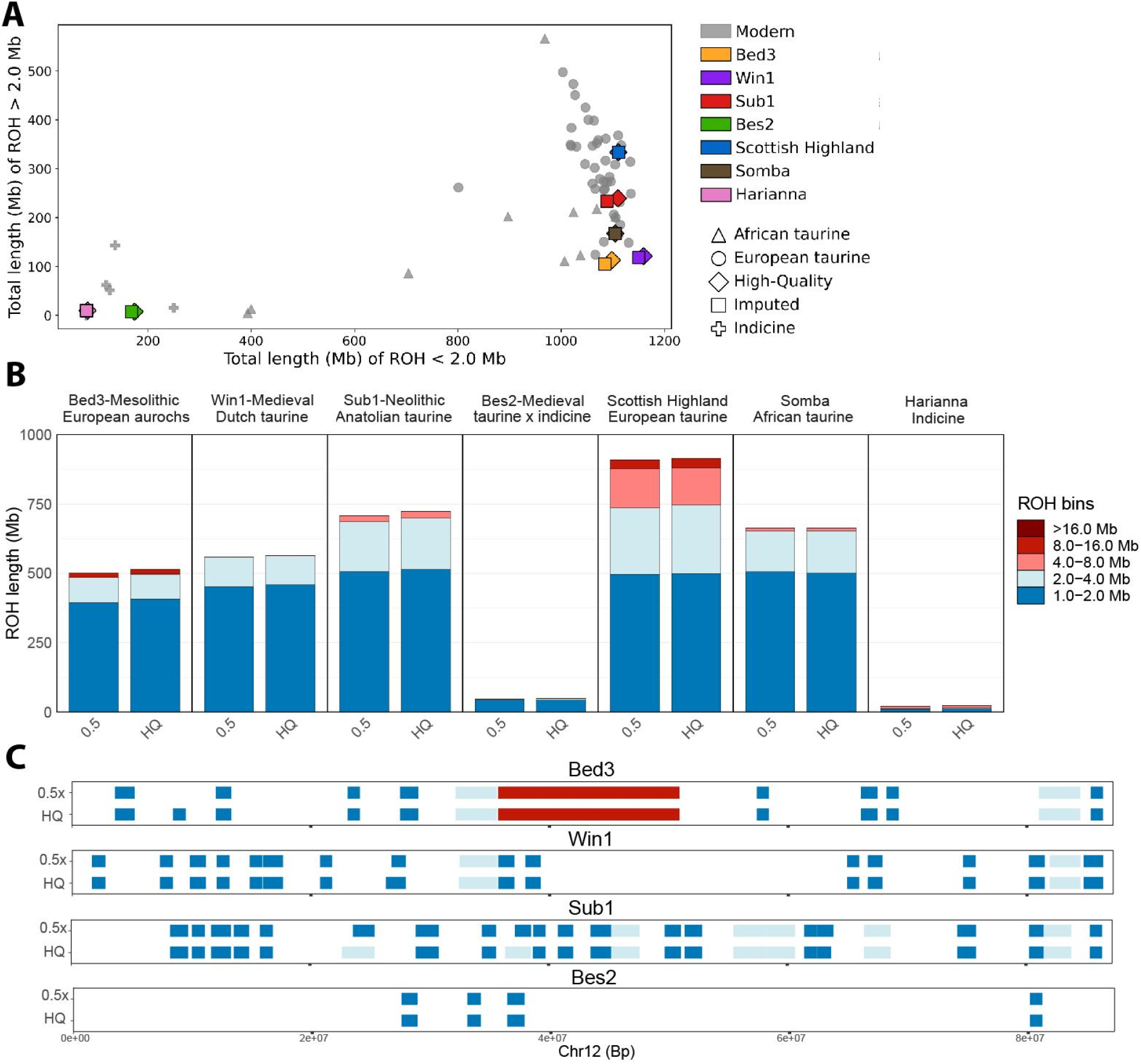
Runs of Homozygosity (ROH) estimates for the 0.5× imputed and the high-quality genotypes. **A**: The total length of ROH >2Mb and ROH ≤2Mb plotted with a subset of the reference panel as a background in grey. Filtering included MAF ≥2.5%, transversions only and no missingness. **B**: The total length of ROH split into length bins for the seven test samples. The number of sites used for analysis = 481,786 filtered for a MAF ≥2.5%, transversions only and no missingness. **C:** Runs of homozygosity along Chr12 for the four ancient genomes at 0.5×, and high-quality demonstrating the general consistency between the imputed and high-quality dataset; filters are the same as B.

Moreover, imputation seems robust for estimating both recent genealogical and deeper population histories via alternately small (<2 Mb) and large ROH (Figure 4A). In the test samples the most pronounced inbreeding is in the modern Scottish highland sample; a feature typical of European production breeds (Figure SI 10) (Purfield et al., 2012) and a pattern absent from the ∼1.5kyr Dutch sample. The ancient zebu hybrid, Bes2, had markedly low levels of ROH, reflecting the large effective population size and genome diversity that is a well-established feature of *Bos indicus* history (Bovine HapMap Consortium et al., 2009; Chen et al., 2018; Murray et al., 2010).

We demonstrate that imputation of ancient cattle, including of the extinct European aurochs, is a feasible methodology for future studies. The success of the imputation of aurochs implies segregating haplotypes in the modern reference panel, most likely from introgression. While damage is potentially disruptive, this is correctable with a deamination aware approach. Imputation accuracy is high and is relatively consistent across the downsampled coverages, demonstrating the feasibility of imputing ancient genomes as low as 0.5× or even lower. This is demonstrated through the consistency in the analysis between imputed and high-quality genotypes. The successful imputation of ancient cattle presents the opportunity for haplotype aware analysis in the future.

## Materials and Methods

### Ancient Processing - Bed3 & Win1

The two archaeological samples were processed in dedicated ancient DNA laboratories in Trinity College Dublin (Win1) and Johannes Gutenberg University of Mainz (Bed3).

#### Win1

A petrous bone, GIA collection number 3848, was excavated in 1997 from the site Winsum-Bruggeburen in the Netherlands. Site occupation was from 7th century BC to 14th century AD, and is thought to have been a military Roman outpost (Bos et al., 1997).

Sample preparation of Win1 was performed as described in (Verdugo et al., 2019). In brief, a wedge of bone was drilled and subsequently powdered using a mixer mill. DNA extraction of ∼150mg bone powder followed a 48 hour 2-step double extraction of two 24 hour digestion steps (37°C) with fresh extraction buffer (0.5M EDTA pH8, 1 M Tris-HCl, 2% sodium dodecyl sulphate, and 100 μg/mL proteinase K) added after the first 24 hour period. Digestion was followed by a Tris-EDTA wash using Amicon® Ultra 4 mL filter (Merck Millipore) followed by a DNA purification step using the “QIAQuick minElute purification kit” (Qiagen) and eluted in 40 μL of EB buffer + Tween® 20 (Sigma-Aldrich) (Mattiangeli, 2023). Prior to library preparation a UDG-treatment was performed using 16.25ul of purified DNA and 5 μl (1U/1uL) USER enzyme (New England BioLabs®, Inc.) and an incubation of 3 hours at 37°C. Double stranded libraries were prepared (Meyer & Kircher, 2010) and sequenced (100 bp SE) on an Illumina Hiseq 2000 at Macrogen, Inc (1002, 254 Beotkkot-ro, Geumcheon-gu, Seoul, 153-781, Republic of Korea).

Radiocarbon dating was performed at 14CHRONO at Queen’s University Belfast (UBA-29049 1556 ± 31 BP) and calibrated dates produced by OxCal 4.4 IntCal20 atmospheric curve (Ramsey, 2009; Reimer et al., 2020). (Figure SI 12)

#### Bed3

A petrous from an aurochs skull, find number 104/102-1, was excavated from the Mesolithic site of Bedburg-Königshoven in Germany (Street, 1999) along with numerous animal bones and two red deer skull headdresses (Wild et al., 2020). The site – today destroyed by lignite mining – was a dump area, formerly located at the bank of a prehistoric wetland close to the former river Erft (Street, 1999, 2020).

Sample preparation of Bed3 was performed as described for sample CTC in (Botigué et al., 2017) and Ch22 and Th7 in (Verdugo et al., 2019), with the following alterations; two independent extractions of 1g of bone powder were performed with a pre-lysis EDTA wash for 30 minutes at room temperature. Extracts underwent UDG-treatment and subsequently fifteen double stranded sequencing libraries (Meyer & Kircher, 2010) were prepared and sequenced (100 bp SE) on an Illumina Hiseq 2000 at Macrogen, Inc (1002, 254 Beotkkot-ro, Geumcheon-gu, Seoul, 153-781, Republic of Korea).

Radiocarbon dating was performed by CologneAMS at the University of Cologne in 2014 as part of the Mesolithic project D4 of the CRC 806 (COL-2680-2.1 - 10036 ± 42) and calibrated dates produced by OxCal 4.4 IntCal20 atmospheric curve (Ramsey, 2009; Reimer et al., 2020) (Figure SI 13)

### Ancient Genome Alignment

Fastq files were processed through a pipeline similar to (Verdugo et al., 2019). Reads were trimmed for adapter sequences using cutadapt (v. 1.1) (Martin, 2011) (*-0 1 & -m 30*) and aligned to ARS-UCD1.2 with the addition of the Y from BosTau5 using the Burrows-Wheeler Alignment (v. 0.7.5a-r405) (Li & Durbin, 2009) with the sub-command aln (*-l 1024 -n 0.01 -o 2*). BAM files were sorted with SAMtools (v. 1.9) (Li et al., 2009) and duplicates removed with Picard (v. 2.20.3) (“Picard Toolkit,” 2019). Indel realignment was performed using the Genome Analysis ToolKit (v. 3.3.0)(McKenna et al., 2010), SAMtools implemented for mapping quality filtering (*-q25*) and coverage calculations performed by Qualimap (v. 2.1.3)(Okonechnikov et al., 2015). To further minimise the effects of deamination, the soft clipping of five base pairs at both ends of the reads was performed.

### Modern Genome Processing

Publically available fastq files of 201 individuals (Table SI 6) were downloaded and processed through the following pipeline. Reads were trimmed for adapters using Trimmomatic (v 0.39) and aligned with BWA mem (v. 07.13)(Li, 2013). Reads were sorted and duplicates removed (PICARD 2.20.3)(“Picard Toolkit,” 2019) and properly paired reads retained (SAMtools v 1.9)(Li et al., 2009). Indel alignment was performed (GATK version v. 3.3.0)(McKenna et al., 2010) and reads filtered via samtools for mapping quality (q 25).

### Variant Discovery

Variants (single nucleotide polymorphisms (SNPs) and insertions or deletions (INDELs)) were called from 201 mapped and filtered bam files of modern cattle (Table SI 6) with an average coverage above 7.6× using Graphtyper (Eggertsson et al., 2017), running each chromosome in parallel (Version 2.7.4). SNPs were removed if they were within 3bp of another INDEL or SNP with vcftools 0.1.17. INDELs were removed and SNPs were were filtered for bi-allelic alleles, a minimum genotype depth of 6×, a maximum genotype depth of 3 times the average genomic coverage of that individual, a minimum quality of 25 and a minimum genotype quality of 20 with vcftools version 0.1.17. SNPs were further filtered according to Graphtyper’s guidelines with bcftools filter version 1.12 *(QD > 2.0, SB < 0.8, MQ > 40.0, LOGF > 0.5, AAScore > 0.5) (Li, 2011)*. As a final filtering step singletons, repetitive regions and genotypes with more than 20% missingness were removed with vcftools version 0.1.17 (Danecek et al., 2011), resulting in 21,656,053 high-quality SNPs, Table SI 2. After filtering, individuals with more than 20% missingness and individuals with 2^nd^ degree relatives (>0.0885 score, vcftools relatedness2) were removed from the dataset resulting in 171 individuals (Table SI 2 & 6). The reference panel consists of 75 European *Bos taurus*, 27 Asian *Bos taurus*, 15 African *Bos taurus*, 10 African *Bos taurus x Bos indicus*, 5 southwest Asian *Bos taurus x Bos indicus*, 23 North East Asian *Bos taurus x Bos indicus* and 16 *Bos indicus*. The reference panel was phased using Beagle5 (Browning & Browning, 2007), with the parameters *impute=false, window=40, overlap=4, gp=true ne=20000*.

### Pseudo-haploid Dataset

The previously published ancient low– to medium–coverage genomes (0.1-3.8x coverage range; Table SI 1) were pseudo-haploidized using ANGSD version 0.938 (Korneliussen et al., 2014) *doHaploCall*, with the following parameters: *doHaploCall 1*, *doCounts* 1, *dumpCounts* 1, minimum base quality of 30 (*-minQ 30*), minimum mapping quality of 25 (*-minMapQ 25*), retain only uniquely mapped reads (*-UniqueOnly 1*), remove reads flagged as bad (*-remove_bads*), remove triallelic sites (*-rmTriallelic 1e—4*), downscale mapping quality of reads with excessive mismatches (*-C 50*), discard 5 bases of both ends of the read (*-trim 5*), remove transitions (-rmTrans 1). The abovementioned sites in the modern reference panel (21,656,053 high-quality SNPs) were used as input for ANGSD using the parameter *–sites*. As a sanity check, the major/minor alleles of the low coverage ancient were compared to the modern reference panel and were removed if there were any discrepancies. ANGSD haplo files were transformed to plink tped files with the haploToPlink function from ANGSD version 0.938 and recoded into ped files with PLINK v.1.90 (Chang et al., 2015). Transitions were removed because transitions, unlike transversions, are most affected by postmortem deamination of DNA, which might increase the number of wrongly called SNPs. The restriction to transversions only is a standard approach in ancient DNA studies.

### Genotype Imputation

Four ancient (13.8-18.7× coverage range) and three modern genomes (28.4-32.7× range) were downsampled to 0.25×, 0.5×, 1.0x and 2.0x on a chromosomal level using picard 2.20.0. The three high coverage (>28x) modern cattle were selected to represent European *Bos taurus* (Scottish Highland - ERR3305587), African *Bos taurus* (Somba - ERR3305591) and Indian *Bos indicus* (Harianna - SRS3120723). The downsampling was performed on chromosomal level so that average genomic coverage would not skew the downsampling process. Genotype likelihoods were computed for the downsampled and the original high-coverage genomes for the high-quality 21,656,053 SNPs mentioned in the Variant calling section.

Genotype calls and likelihoods were generated according to the GLIMPSE version 1.1.1 pipeline (Rubinacci et al., 2021), with the command bcftools mpileup (version 1.12) with parameters *-I,-E, -a “FORMAT/DP,FORMAT/AD,INFO/AD”*, the reference genome and the abovementioned sites in the reference panel (*-T*) followed by bcftools call with the parameters *-Aim -C alleles*, and the abovementioned sites (*-T*). This step was performed on both downsampled and high-coverage genomes. The high-coverage genotype likelihoods were further filtered for a minimum base quality of 30, a minimum genotype quality of 25, a minimum genotype coverage of 8, a maximum genotype coverage of 3 times the average genomic coverage and a minimum allelic balance of 40%, obtaining the validation golden standard genotype likelihoods.

Imputation was performed on the downsampled genomes using GLIMPSE v1.1.1 (Rubinacci et al., 2021), according to the GLIMPSE pipeline. Chromosomes were split into chunks of 2 Mb with a 200kb buffer window with *GLIMPSE_chunk*. Imputation was performed on these chunks with default parameters using *GLIMPSE_phase* with the reference panel created in the section Variant calling. The imputed chunks were ligated using *GLIMPSE_ligate* with default parameters. The imputed data was filtered to keep only the most confidently imputed SNPs, this was done by filtering for a strict genotype probability *(GP) ≥0.99* and an *INFO score ≥0.99*, each sample was imputed and filtered separately. For imputation of the three modern test samples, reference panels without the test individual were created and subsequently used for imputation of the respective test sample.

### Accuracy of Genotype Imputation

Imputation accuracy, seen as genotype concordance between the imputed and the high-quality validation genotypes, was calculated with Picard GenotypeConcordance version 2.20.0 (“Picard Toolkit,” 2019). Imputation accuracy was calculated for heterozygotes, homozygotes alternative and the combined alternative alleles; this was done for eight MAF bins and a MAF threshold. Picards’ GenotypeConcordance tool was used with the high-quality validation genotypes as the *TRUTH_VCF*, the imputed genotypes as the *CALL_VCF*, and for the specific MAF bins mentioned previously with the *INTERVALS* parameter. Genotype concordance, called sensitivity in picards’ output, is calculated as *TP* / (*TP* + *FN*), where TP (true positives) stands for variants where the CALL matches the TRUTH, and FN (false negatives) stands for when variants do not match the reference. Recovery of genotypes is calculated as (*TP* + *FN*)/ *Total HQ*, where Total HQ stands for the genotypes present in the high-quality validation TRUTH. Non-reference-discordance (NRD), a measurement of error rate was calculated following the formula published by (Sousa da Mota et al., 2023), NRD does not take correctly imputed homozygous reference sites into account, giving more weight to imputation errors at alternative sites.

Win1, an European taurine animal, demonstrates the lowest imputation accuracy of heterozygote and homozygote alternative alleles (Figure 2). While all ancient samples underwent UDG-treatment, deamination is still detected at CpG sites and in Win1 this is elevated throughout the first and last 30bp of the sequencing reads (Figure SI 2). We find that in this sample the imputation accuracy can be improved when we filter for potential deamination signals prior to imputation. This was achieved by a) setting heterozygote or homozygote reference genotypes as missing if the reference allele was T or A and the alternate C or G respectively b) setting heterozygote or homozygote reference genotypes as missing if the reference was T or A and the alternate allele was C or G respectively. Using these filtered genotypes we demonstrate an improvement in post imputation accuracy (Figure SI 3).

### Downstream analyses

The filtered imputed and high-quality genotypes were merged with the modern reference panel using PLINK v.1.90 (Chang et al., 2015). In the case of PCA and admixture analyses, the data was filtered for MAF >2.5%, transversions only and linkage disequilibrium (indep-pairwise 50 5 0.5), resulting in 812,801 SNPs. For the ROH analysis, a subset of modern samples <5% missingness (N=60) was created and merged with the imputed genotypes. The filters on the ROH dataset consisted of MAF >2.5%, no genotype missingness and transversions only, resulting in 481,786 SNPs.

#### PCA including ancient pseudo-haploid

The pseudo-haploid ancient samples were merged with the dataset containing the imputed and high-quality genotypes and modern reference panel, filtered for MAF >2.5%, transversions only and linkage disequilibrium (indep-pairwise 50 5 0.5), resulting in 812,801 SNPs. Smartpca version 16000 was used to perform a PCA with default parameters (Patterson et al., 2006; Price et al., 2006). The first 10 principal components were calculated using the modern reference panel, the pseudo-haploid, imputed and high-quality genotypes were projected (*lsqproject:yes)*.

#### PCA to test imputation genotypes

Smartpca version 16000 was used to perform a PCA with default parameters (Patterson et al., 2006; Price et al., 2006). The first 10 principal components were calculated using the modern reference panel, both the imputed and high-quality genotypes were projected (*lsqproject:yes)*. PCA was performed on each imputed coverage separately.

#### Model Based Admixture

ADMIXTURE v1.3.0 (Alexander et al., 2009) was used to estimate ancestry proportions for the modern reference panel, high-quality and imputed genotypes. ADMIXTURE ran for K between two and ten, for the best K’s (4-5), a bootstrap with 1000 replicates (--B 1000) was run to obtain the standard error and bias of admixture estimates. Admixture ran on each imputed coverage separately.

#### Runs of Homozygosity (ROH)

ROH were estimated with PLINK v1.90 (Chang et al., 2015) with the parameters *--homozyg - -homozyg-density 50 --homozyg-gap 100 --homozyg-kb 500 --homozyg-snp 50 --homozyg-window-het 1 --homozyg-window-snp 50 --homozyg-window-threshold 0.05*, according to earlier studies (Cassidy et al., 2016; Gamba et al., 2014; Martiniano et al., 2017; Sousa da Mota et al., 2023). ROH were estimated using a subset of moderns (<5% missingness), high-quality and imputed genotypes for each imputed coverage separately.

## Data Availability

The raw reads for the two new ancient genomes will be deposited with ENA (Project XXXX). The accession numbers for the publicly available genomes used in the reference panel are noted in Table SI 1-2 & Table SI 6. The VCF of the phased reference panel will be made publicly available xxxx.

## Acknowledgments

We are grateful to Daniel Bradley for his valuable mentorship, discussions and reading of the paper. We thank Daan Raemaekers, Canan Çakirlar, Joachim Burger and Conor Rossi for their reading of the paper and helpful suggestions. We thank GIA for access to the WIN1 sample. This work has been supported by The European Research Council under the European Union’s Horizon 2020 research and innovation programme (885729-AncestralWeave, 295729-CodeX), Government of Ireland Postdoctoral Fellowship (GOIPD/2020/605) and Dutch Research Council Open Competition (Grant No. 406.18.HW.026).

