## Supplemental Figures for "A high coverage Mesolithic aurochs genome and effective leveraging of ancient cattle genomes using whole genome imputation"

### Supplementary Materials

#### Figures

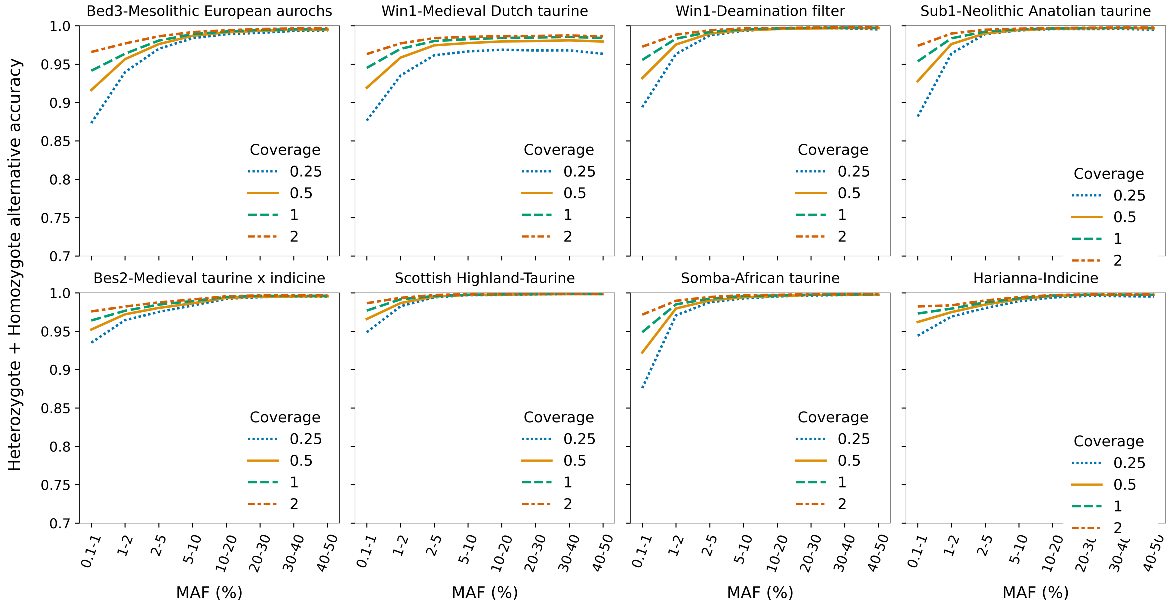


**Figure S1. Accuracy of imputation of heterozygous and homozygous alternative sites at different minor allele frequency (MAF) bins and different downsampled coverages, for all sites.** Downsampled coverage is denoted by line style and colour. An additional graph is included for Win1 to demonstrate the positive effect on the accuracy of a deamination filter prior to imputation.

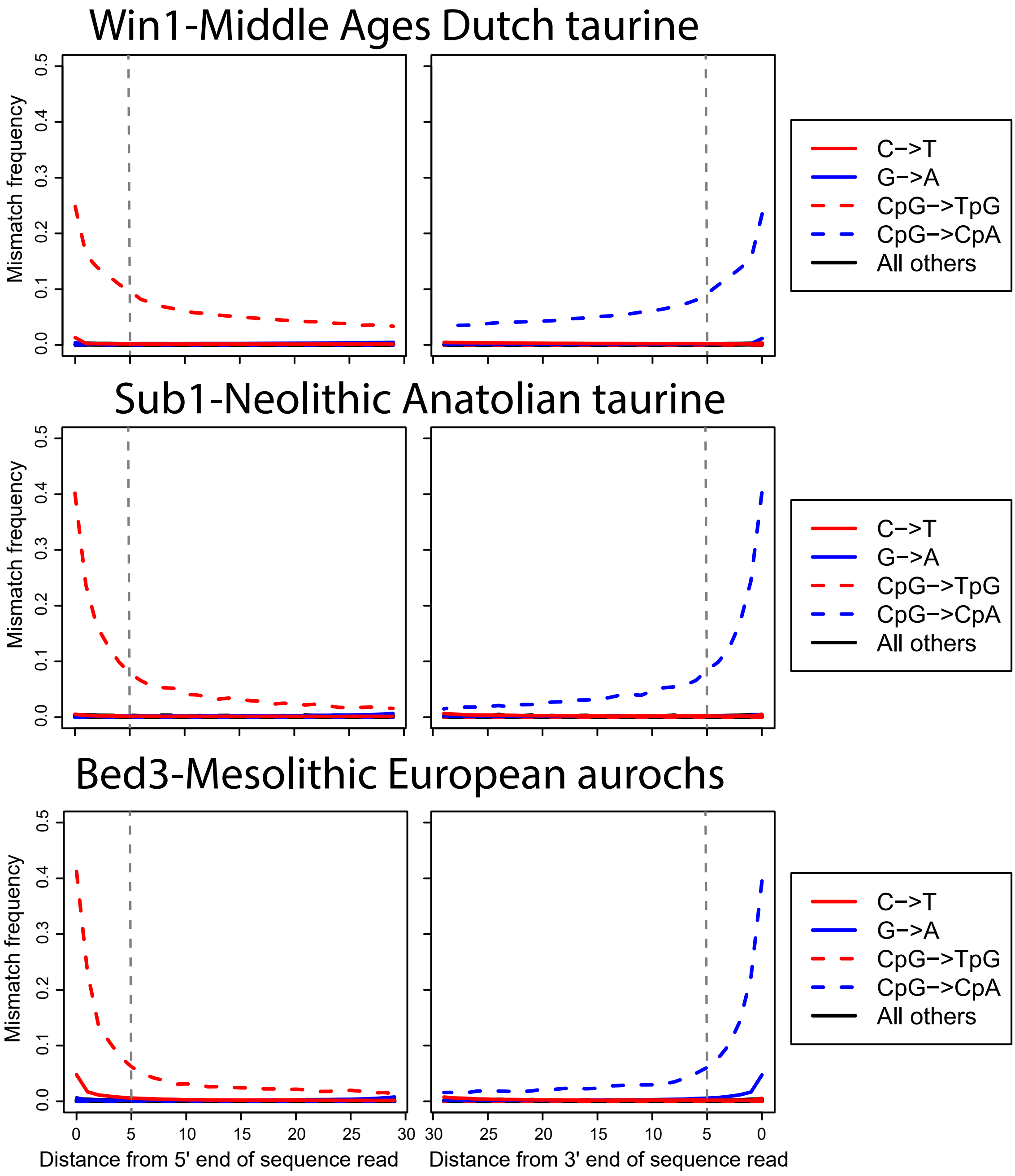


**Figure S2. Damage profiles at CpG sites for Win1, Sub1, Bed3 demonstrating the damage patterns typical of ancient DNA.** This was calculated prior to the soft-clipping of five base pairs from both ends, clipping is denoted by the black dashed line. The soft-clipped BAM files were the input for the imputation pipeline.

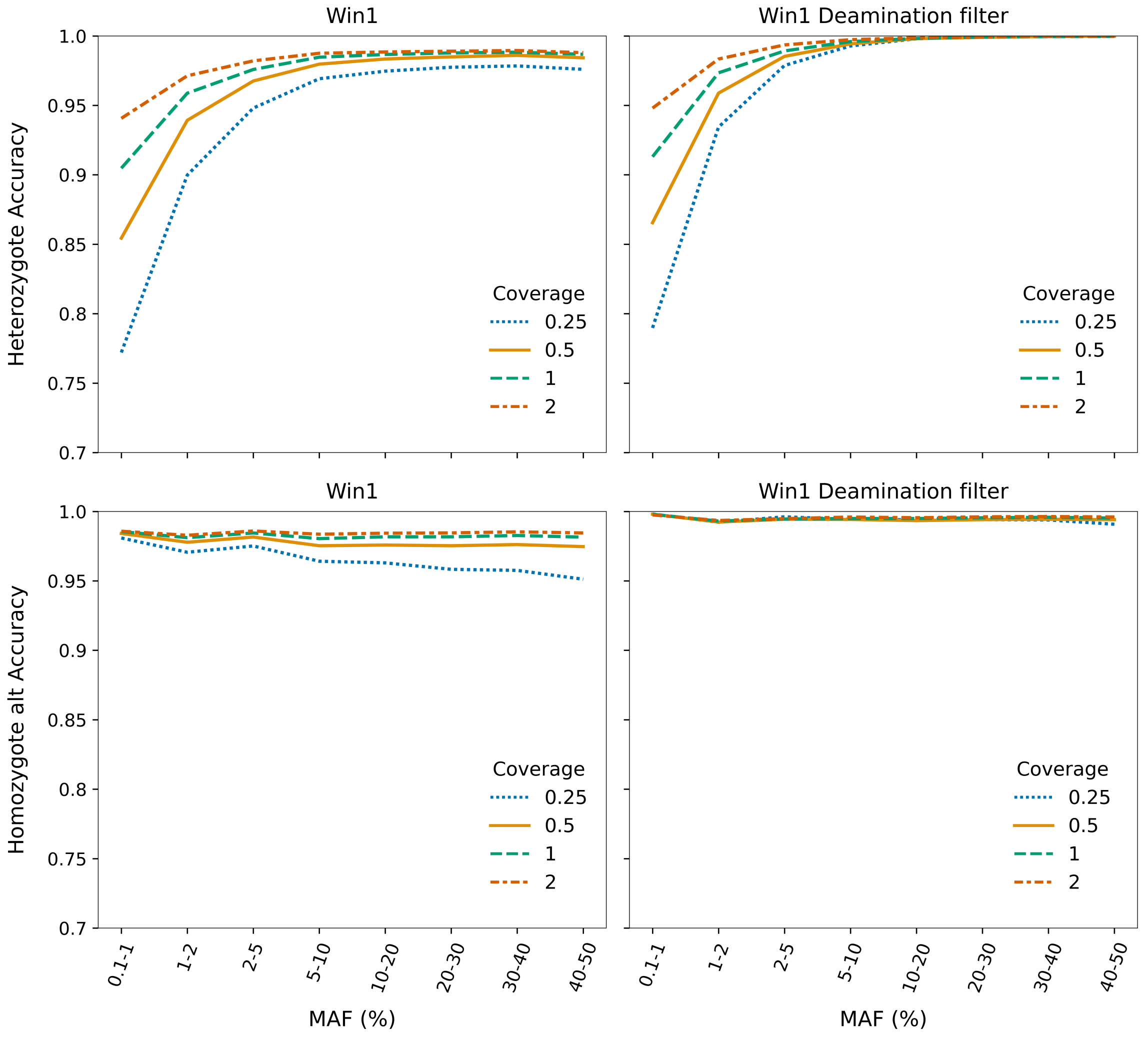


**Figure S3. The imputation accuracy for Win1 without a deamination filter and with deamination filter prior to imputation.** An improvement in accuracy is shown when Win1 was filtered for deamination.

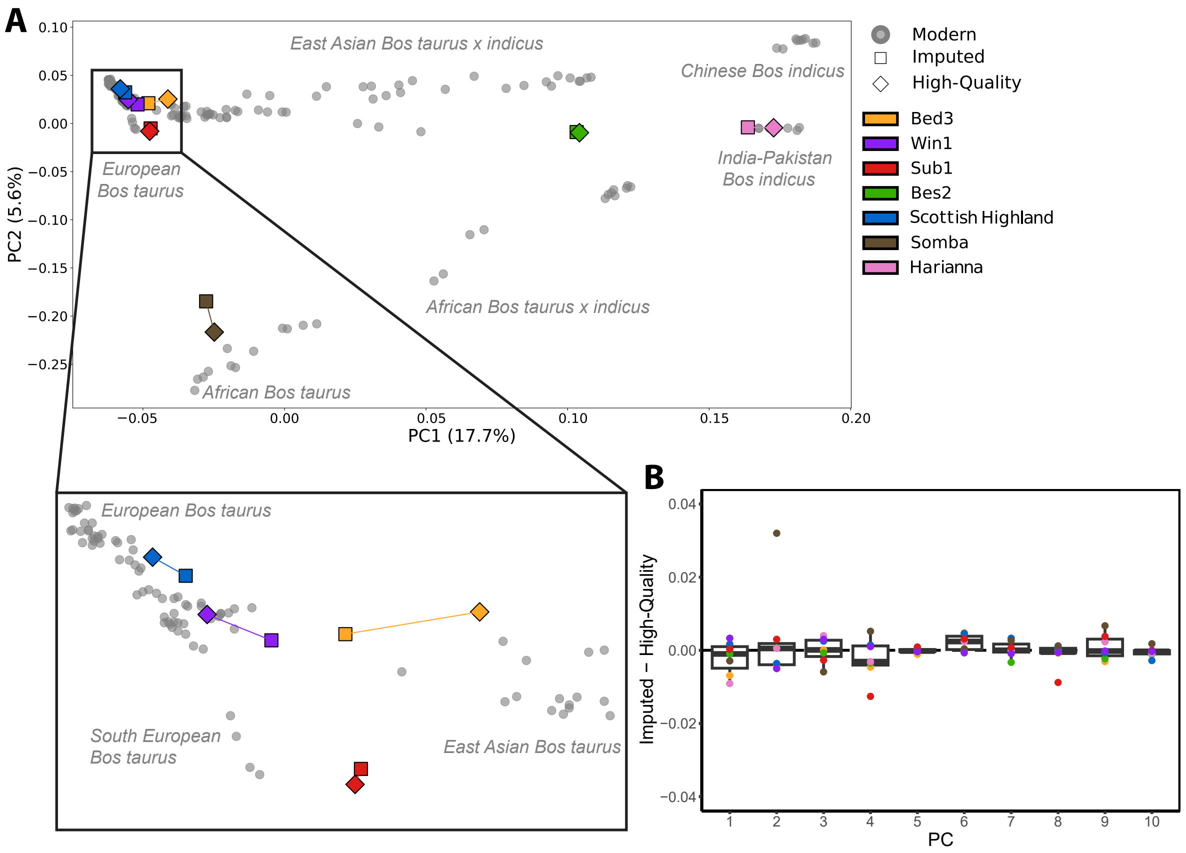


**Figure S4. Principal Components Analysis (PCA) of 0.25x imputed genotypes and high-quality genotypes of the seven test individuals onto the modern cattle reference panel A:** Projections for 0.25x imputed and high-quality genotypes of the test samples onto the modern reference panel along the first two eigenvectors. Dataset was filtered for a MAF ≥2.5%, transversions and LD pruning resulting in 800,694 SNPs. Test individuals are denoted by colour with imputed and high-quality represented by squares and diamonds respectively, while the reference panel individuals are plotted as grey circles. **B:** Boxplots of the normalised differences in the coordinates of the high-quality and 0.25x imputed genomes for the first ten principle components. The horizontal lines of the box pot represent the first quartile, median and third quartile, whiskers represent 1.5 times the quartile range

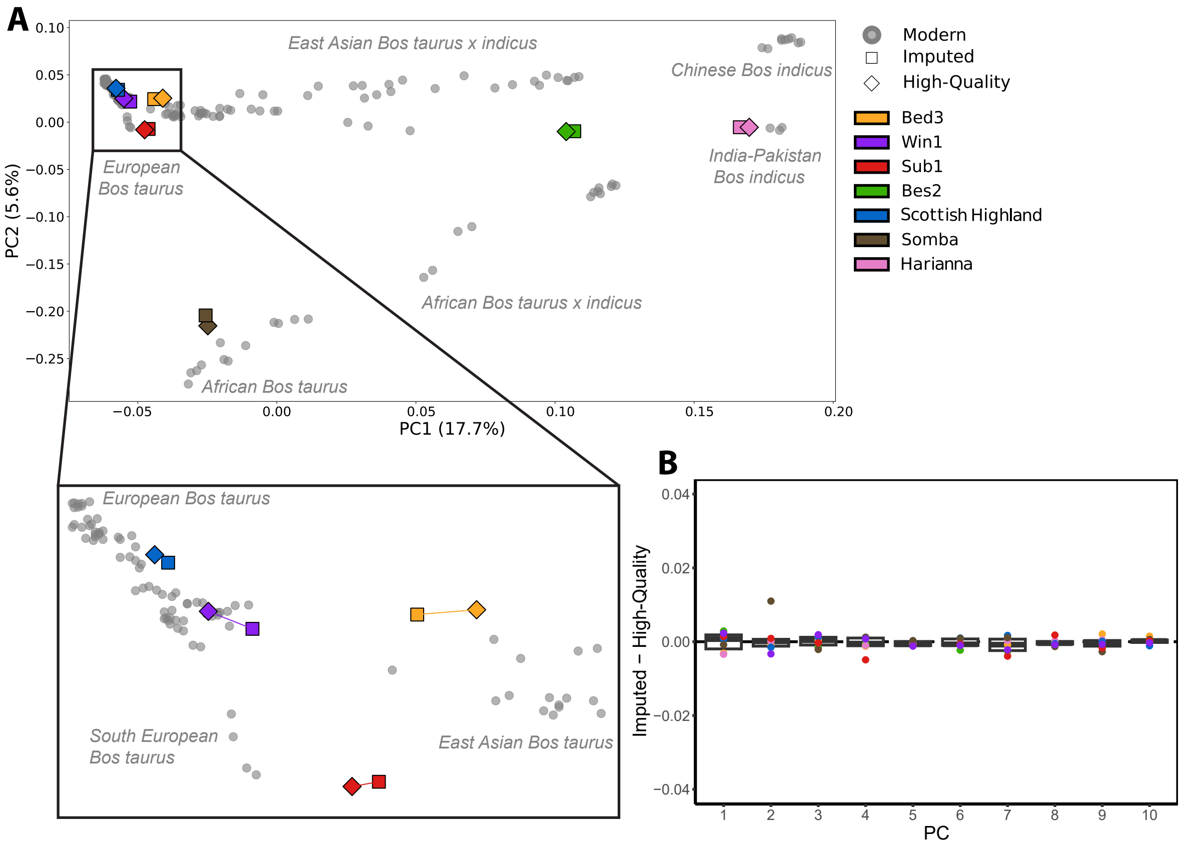


**Figure S4. Principal Components Analysis (PCA) of 1x imputed genotypes and high-quality genotypes of the seven test individuals onto the modern cattle reference panel A:** Projections for 1x imputed and high-quality genotypes of the test samples onto the modern reference panel along the first two eigenvectors. Dataset was filtered for a MAF ≥2.5%, transversions and LD pruning resulting in 840,555 SNPs. Test individuals are denoted by colour with imputed and high-quality represented by squares and diamonds respectively, while the reference panel individuals are plotted as grey circles. **B:** Boxplots of the normalised differences in the coordinates of the high-quality and 1x imputed genomes for the first ten principle components. The horizontal lines of the box pot represent the first quartile, median and third quartile, whiskers represent 1.5 times the quartile range

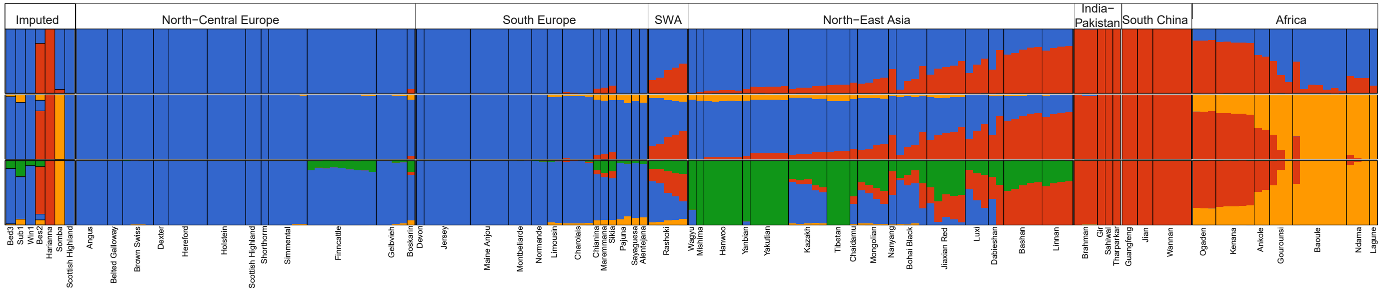


**Figure S6. Model based estimates of ancestral coefficients for the seven 0.5x imputed samples and the reference panel for K2-K4.**
ADMIXTURE analysis based on 800,694 SNPs (MAF ≥2.5%, transversions and LD pruning). Ancestral components are shown for taurine (blue), indicine (red), African taurine (orange) and East Asian taurine (green).
K3-4 had the lowest CV score, and are the optimal K.

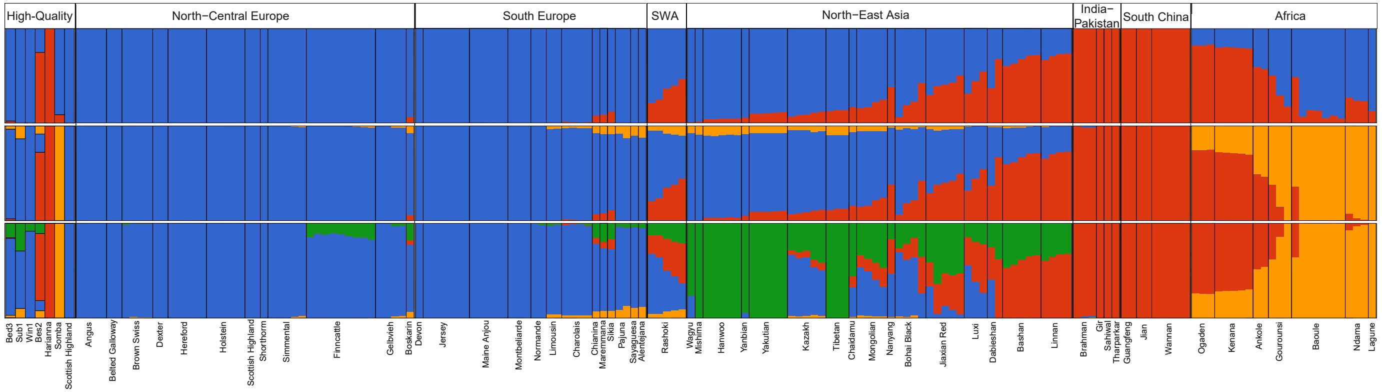


**Figure S7. Model based estimates of ancestral coefficients for the seven high-quality genotypes and the reference panel for K2-K4.**
ADMIXTURE analysis based on 800,694 SNPs (MAF ≥2.5%, transversions and LD pruning). Ancestral components are shown for taurine (blue), indicine (red), African taurine (orange) and East Asian taurine (green).
K3-4 had the lowest CV score, and are the optimal K.

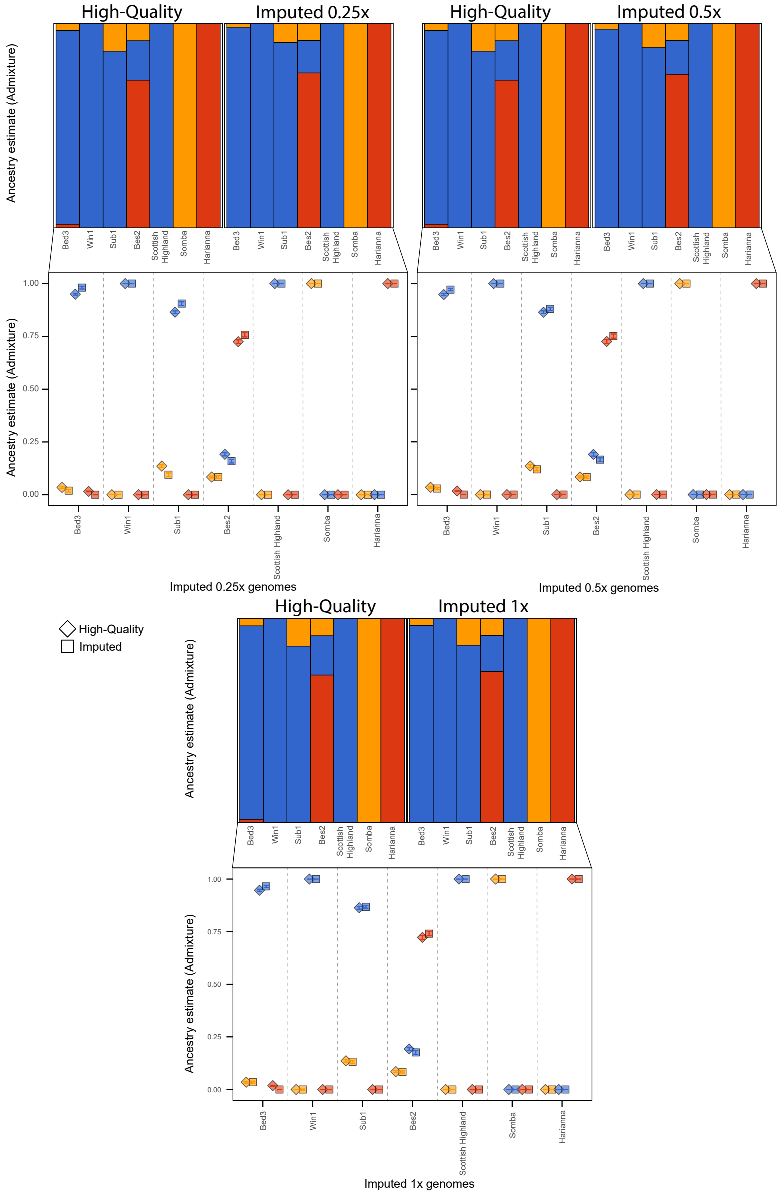


**Figure S8. Comparison of the model based estimates of ancestral coefficients for the high quality genotypes and imputed genotypes for K=3.** ADMIXTURE analysis based on 800,694 SNPs (MAF ≥2.5%, transversions and LD pruning) run with the reference panel and the seven test samples, only the test samples are plotted for ease. Ancestral components for K=3 are displayed for the 0.25x, 0.5x and 1x imputed (squares) and high-quality validation (diamonds) genotypes of the seven samples, colours are denoted as taurine (blue), indicine (red) and African taurine (orange). Error bars represent the deviation from the mean measured as standard errors, calculated by bootstrapping 1000 times.

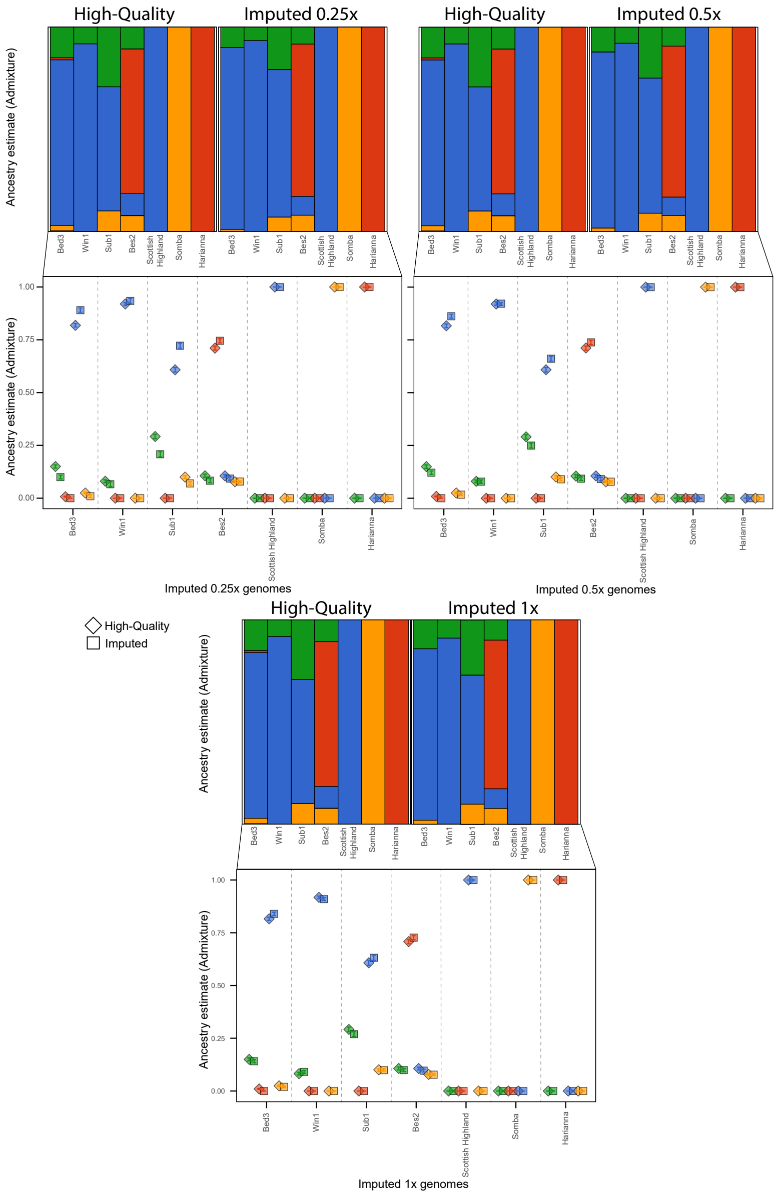


**Figure S8. Comparison of the model based estimates of ancestral coefficients for the high quality genotypes and imputed genotypes for K=4.** ADMIXTURE analysis based on 800,694 SNPs (MAF ≥2.5%, transversions and LD pruning) run with the reference panel and the seven test samples, only the test samples are plotted for ease. Ancestral components for K=4 are displayed for the 0.25x, 0.5x and 1x imputed (squares) and high-quality validation (diamonds) genotypes of the seven samples, colours are denoted as taurine (blue), indicine (red) and African taurine (orange). Error bars represent the deviation from the mean measured as standard errors, calculated by bootstrapping 1000 times.

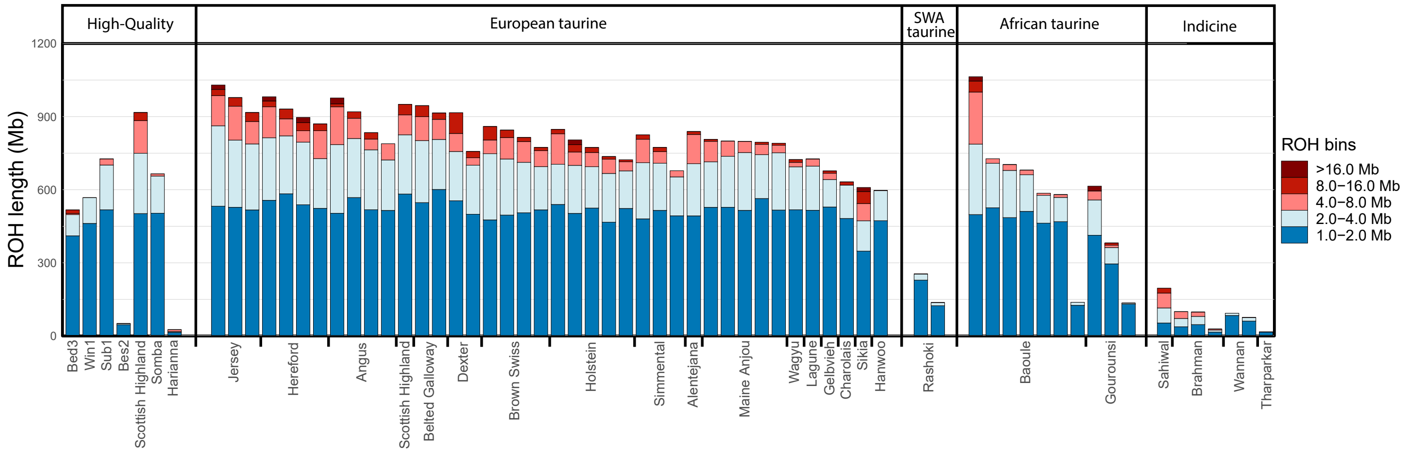


**Figure S10. Runs of Homozygosity (ROH) estimates for the high-quality genotypes and the modern reference panel.** The total length of ROH split into length bins for the seven test samples and a subset of the modern reference panel (N=60). The number of sites used for analysis = 481,786 filtered for a MAF ≥2.5%, transversions only and no missingness.

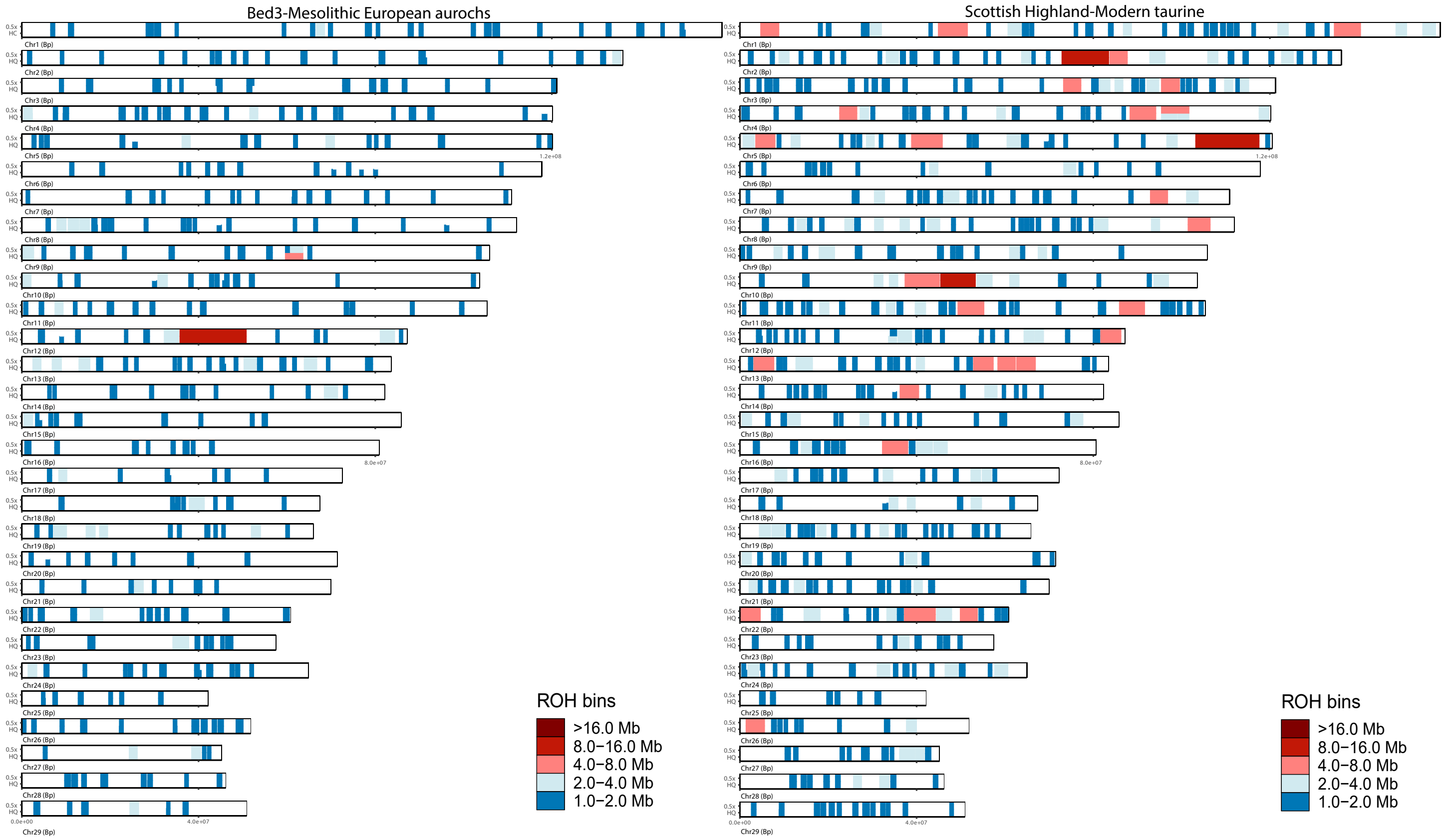


**Figure S11. Runs of homozygosity along all chromosomes for Bed3 and the modern Scottish Highland for high-quality genotypes and imputed genotypes from the 0.5x downsample.** The number of sites used for analysis = 481,786 filtered for a MAF ≥ 2.5%, transversions only and no missingness.

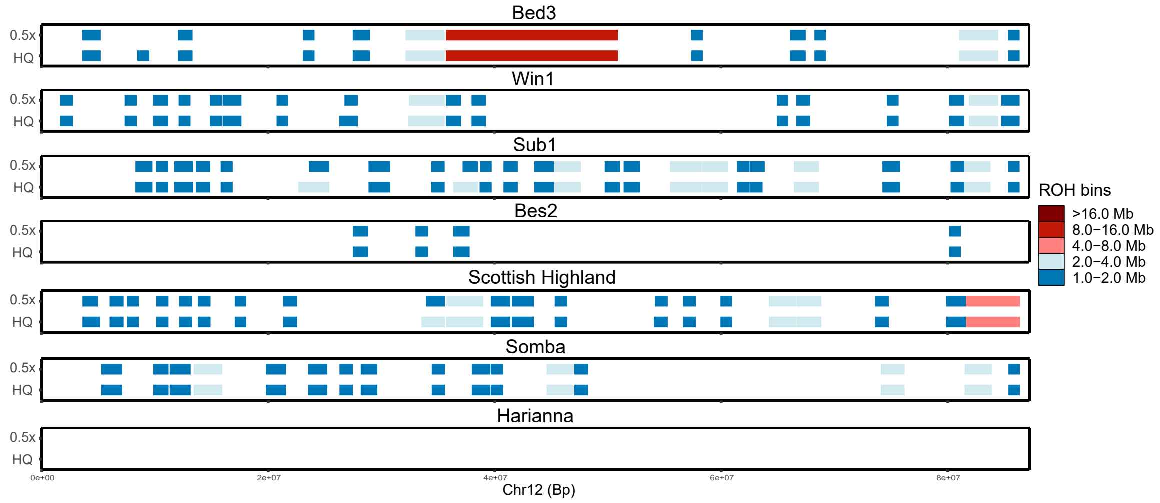


**Figure S12. Runs of homozygosity along chromosome 12 for the seven test samples at 0.5x imputed, and high-quality genotypes demonstrating the general consistency between the imputed and high-quality dataset.** The number of sites used for analysis = 481,786 filtered for a MAF ≥ 2.5%, transversions only and genotype missingness.

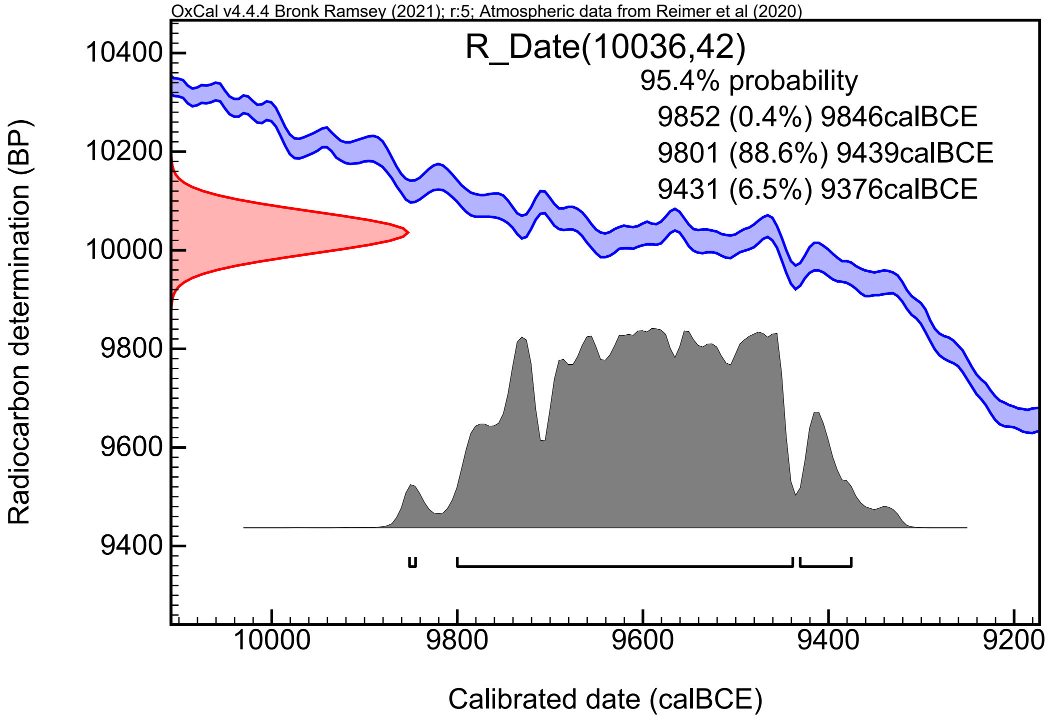


**Figure S12. Radiocarbon calibration curve for Bed3**

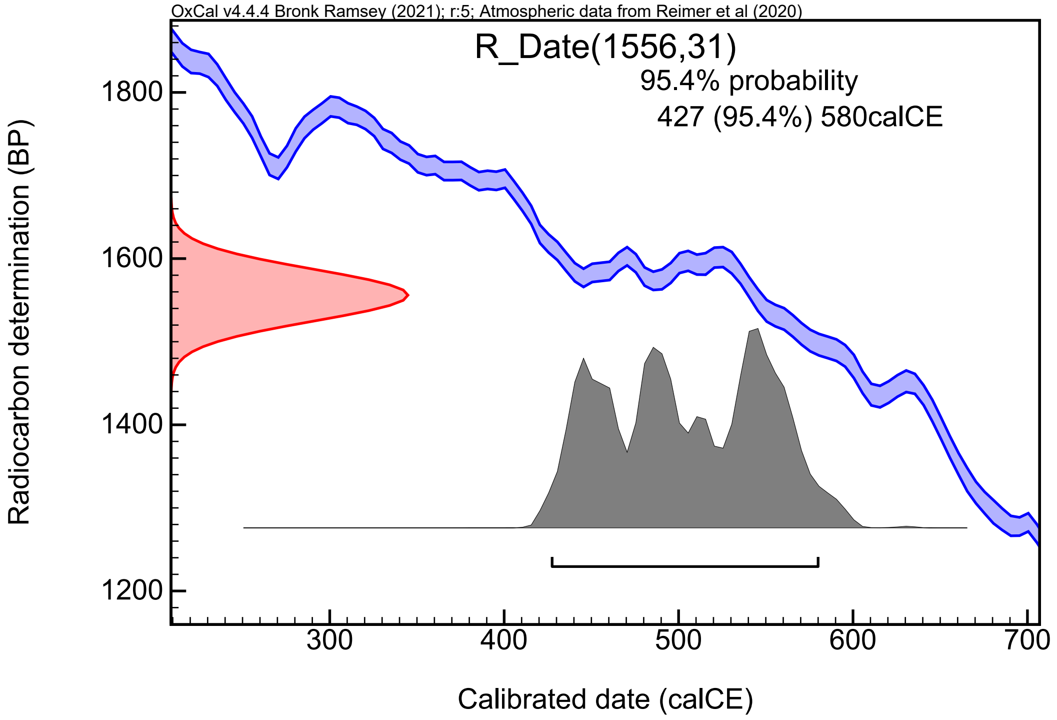


**Figure S12. Radiocarbon calibration curve for Win1**
